# Topologically Optimized Intrinsic Brain Networks

**DOI:** 10.1101/2025.02.19.639110

**Authors:** Noah Lewis, Armin Iraji, Robyn Miller, Oktay Agcaoglu, Vince Calhoun

## Abstract

The estimation of brain networks is instrumental in quantifying and evaluating brain function. Nevertheless, achieving precise estimations of subject-level networks has proven to be a formidable task. In response to this challenge, researchers have developed group-inference frameworks that leverage robust group-level estimations as a common reference point to infer corresponding subject-level networks. Generally, existing approaches either leverage the common reference as a strict, voxel-wise spatial constraint (i.e., strong constraints at the voxel level) or impose no constraints. Here, we propose a targeted approach that harnesses the topological information of group-level networks to encode a high-level representation of spatial properties to be used as constraints, which we refer to as Topologically Optimized Intrinsic Brain Networks (TOIBN). Consequently, our method inherits the significant advantages of constraint-based approaches, such as enhancing estimation efficacy in noisy data or small sample sizes. On the other hand, our method provides a softer constraint than voxel-wise penalties, which can result in the loss of individual variation, increased susceptibility to model biases, and potentially missing important subject-specific information. Our analyses show that the subject maps from our method are less noisy and true to the group networks while promoting subject variability that can be lost from strict constraints. We also find that the topological properties resulting from the TOIBN maps are more expressive of differences between individuals with schizophrenia and controls in the default mode, subcortical, and visual networks.

## 1. Introduction

In the quest to estimate data-driven subject-level functional brain networks in a quantifiable way, group-inference frameworks [1] have become prime contenders. These frameworks primarily use reference networks from methods such as group-independent component analysis (gICA) to estimate individual subject networks at the static [2] and/or dynamic [3] level by leveraging reference networks from either the group-level or pre-computed templates [4]. For example, spatio-temporal regression (STR) [5, 6] uses ordinary least squares (OLS) to estimate the subject-level network timecourses from the reference maps and, subsequently, the subject-level spatial maps from these timecourses. Other frameworks, such as spatially-constrained ICA [7, 8], and template ICA [9] use subject-specific ICA with spatial constraints between a given subject and the reference maps at the voxel level for maps that are reflective of a “reference” image (e.g., the reference-level or template maps). Spatially constrained ICA allows for network estimation from short data lengths [10, 11], but such constraints may limit extensive descriptions of individual-specific properties. The ultimate goal of subject-level network estimation is to provide subject maps that are reflective of the individualized expression of a reference-level network, hence allowing correspondence among subjects, while also preserving individual differences.

The estimation of corresponding subject-level networks across multiple subjects is a difficult problem, which has been generally resolved using reference-level information [6, 12]. However, most methods only consider certain properties of the reference maps, e.g., the statistical or voxel-level information [2, 4]. Instead, we argue that incorporating the geometric properties of the reference maps, and the topology in particular, will alleviate many of the aforementioned limitations. Not only are global geometric properties useful for network map estimation, but they can also provide a wealth of information about the brain as a whole [13].

Existing frameworks for independence-based subject network estimation (e.g., ICA) come with some trade-offs. For example, subject-level ICA provides subject-level networks that are essentially blind. While flexible, this makes identifying correspondence to existing networks or across subjects challenging. Post-hoc methods, such as STR, are straightforward to implement, but treat the ICA solution as fixed weighted maps, ignoring indepen-dence among networks when applied to other data. Conversely, subject-level, spatially constrained ICA estimation methods adapt to individual subjects and leverage higher-order statistics but can still reduce subject-level information and variation in the spatial maps between subjects. All of these approaches ignore certain reference-level information, such as global and local geometries.

The use of topological information and topological data analysis (TDA) is becoming widely used in neuroimaging research. Some of the earliest work leveraged topological information to improve surface reconstruction and segmentation [14, 15]. New methodologies have tried to explain the brain with topological analysis, most relying on graph representations of fMRI data [16, 17]. In addition, some work leverages topological properties to find novel and useful representations of fMRI dynamics [18]. Beyond the field of neuroimaging, there is now burgeoning work in ways to improve machine learning methods with topological information [19]. Based on this, our preliminary work improved subject-level estimation for 2D slices of intrinsic connectivity network (ICN) maps [20], which, to our knowledge, was the first time topological data analysis has been used for network estimation. In this work, we extend that methodology to the entire brain, in three dimensions, and include a much larger feature space of simplicial complexes.

Our proposed method focuses on leveraging topological properties to improve subject-level estimates, building from a previously described method that adds a topological layer for machine learning models [19]. Section 2.4.2 describes their method in more detail. For this purpose, instead of voxel-wise constraints, we impose a constraint on the topological information of global brain networks. Constraining the subject-level linear estimation to be topologically similar to the reference-level maps. Using STR, we first compute the temporal estimations from the reference maps and then compute the spatial estimations with the additional topological loss, which essentially gives us two loss terms in one optimization function: the linear estimation loss and the topological constraint. To implement this topological constraint, we characterize the topologies of the two maps using the persistence diagrams (PDs) of the maps [21, 22]. These PDs are an estimation of the topological properties of the maps, i.e., the number of connected components and the number of holes. Given these PDs for the reference and subject-level maps, our loss function is then defined as the Wasserstein distance between the two PDs, which is theoretically defined as the optimal transport loss in persistence space [21]. This provides a ‘soft’ constraint, while also encouraging correspondence among subjects.

To show that this methodology represents an effective improvement, we compare it to the OLS-only method along three primary axes. Although encouraging the subject maps to be more similar to the reference maps may improve the image quality, we are concerned with negative impacts on the subject maps, namely reduced subject variation. Firstly, as a sanity check, we show that the TOIBN maps are both topologically and statistically more similar to the reference maps than the OLS-only method. Secondly, we show that the topological correction promotes subject variability, primarily in areas that are most relevant to the reference maps. These results attempt to alleviate any concerns that our method is removing important subject-level information. Thirdly, we show that the TOIBN maps are significantly less noisy than the OLS-only maps by measuring the contrast-to-noise ratio (CNR) of each map. This represents the core of the benefits of our method. Finally, we show that the topological correction produces maps that reveal more topological group differences between individuals with schizophrenia and controls. Here, we show again that the methodology improves subject-level information in the form of group differences, while also showing that the topological properties can be useful measures.

## 2. Methods

### 2.1. Data and Preprocessing

For this work, we chose a dataset of fMRI resting-state, eyes-closed images from 182 individuals with schizophrenia and 176 controls for a total of 358 subjects from the Function Biomedical Informatics Research Network (fBIRN) [23]. Each sample has 162 volumes with a TR of 2. The scans come from 7 sites, 6 of which used a 3T Siemens Tim Trio scanner, while the 7*^th^* site used a 3T GE Discovery MR750. The acquisition used gradient-echo echo planar imaging, an FOV of 220 x 220, TE = 30ms, FA = 770, and 32 sequential ascending axial slices of 4mm thickness and 1mm skip.

The images were preprocessed with the Statistical Parametric and Mapping (SPM) (www.fil.ion.ucl.ac.uk/spm/) and Analysis of Functional NeuroImages (AFNI) (afni.nimh.nih.gov/) toolboxes. Motion correction was performed with the INRIAlign toolbox. Slice-timing correction was performed using the middle slice as the reference. Despiking, warping to the Montreal Neurological Institute (MNI) template, resampling to 3mm^3^ voxels, spatial smoothing with a 6mm full-width/half-max Gaussian kernel, and z-scoring were also performed.

### 2.2. Template ICA

We use Neuromark, a template of ICA components based on group ICA as our reference maps [4, 10, 24]. Given a large set of samples across multiple datasets, this pipeline starts with subject-level principal component analysis (PCA) to select the principal components (PCs) with a variance greater than 99%. Then, group-level PCA was applied to the subject-level PCs concatenated across time to select the 20 group-level PCs with the highest variance. The infomax ICA algorithm [25] was applied using the ICASSO framework for 100 runs to obtain the 20 most stable independent components. We selected the components with their peak overlap with the gray matter and the components where low-frequency fluctuations dominate the timecourses. Finally, we found the ones that best visually matched known networks, leaving us with 14 final networks as our reference maps.

### 2.3. Subject-level ICNs Through Spatiotemporal Regression

Reference spatial maps (examples in figure 1) of ICNs have been vital for neuroimaging research. However, it is also imperative to find methods that estimate these networks at the subject level. One approach, spatio-temporal regression (STR) [5, 26], uses linear regression to estimate the subject ICN timeseries and a secondary regression that estimates the subject spatial maps from the computed timeseries. The timecourse estimation is computed as 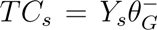, where *Y_s_* is the subject’s fMRI scan, and *θ_G_* is the reference network. From this, the subject network is estimated by 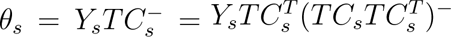 *θ_s_* and *TC_s_* are the subject network spatial maps and timecourses, respectively.

**Figure 1:**
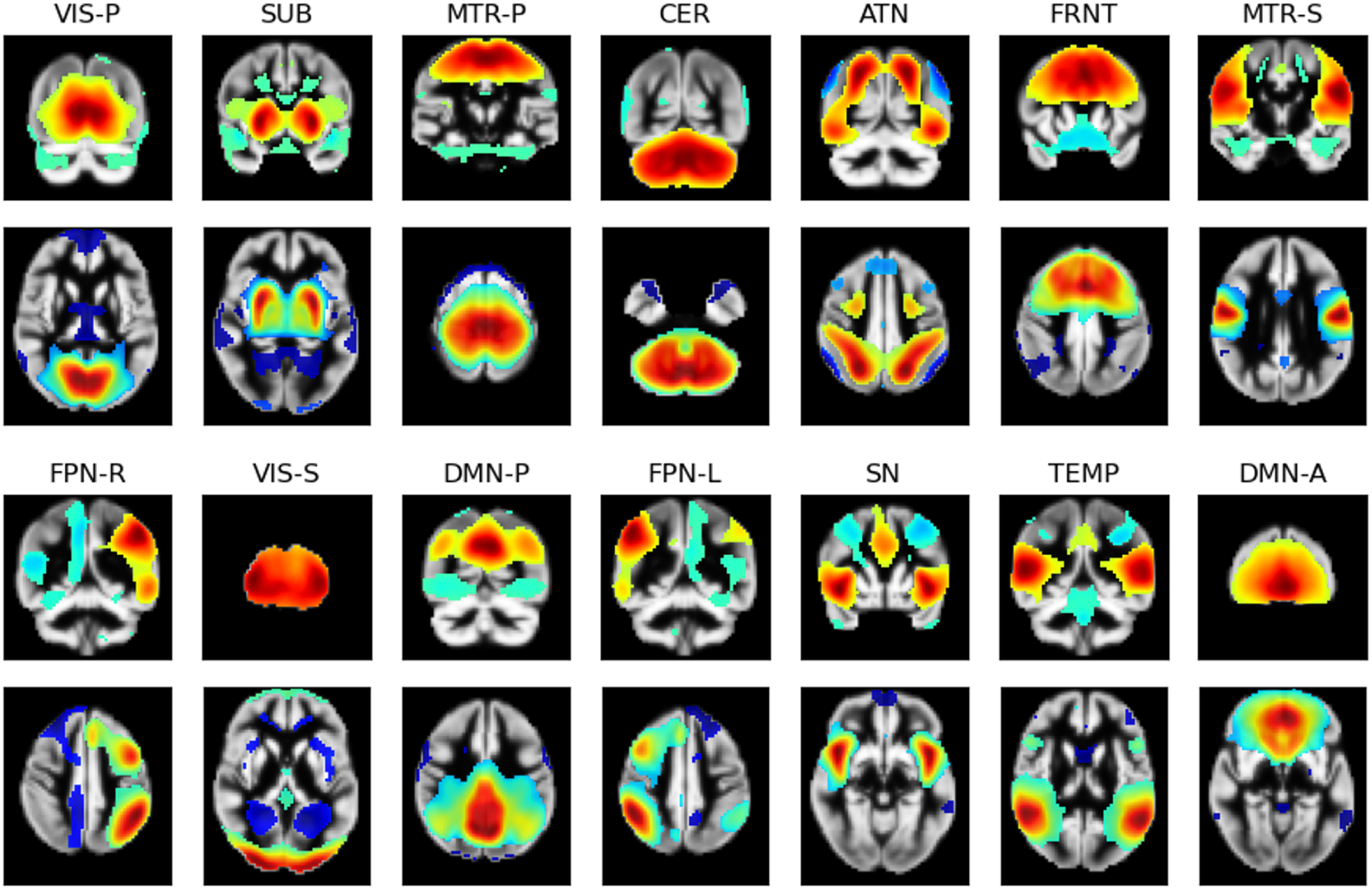
The reference maps, thresholded to the top 30% of absolute-valued voxels, retaining only the highest-valued (red) and lowest-valued (blue) parts of the networks. The networks are defined as: primary visual (VIS-P), subcortical (SUB), primary somatomotor (MTR-P), cerebellum (CER), attention (ATN), frontal (FRNT), secondary somatomotor (MTR-S), frontoparietal-right (FPN-R), secondary visual (VIS-S), posterior default mode (DMN-P), frontoparietal-left (FPN-L), salience (SN), temporal (TEMP), and anterior DMN (DMN-A).

### 2.4. Topology

Topological principles have been an important part of surface reconstruction for decades. Topology is the study of the properties of mathematical objects that are preserved under continuous deformations, such as twisting, stretching, and bending [22]. More simply, these deformations do not close holes, open holes, tear, glue, or cause self-intersections of an object. These continuous deformations are known as “homeomorphisms”. Fundamentally, topology defines the most basic space, a “topological space”, as a set, *X*, and a topology 𝒯. The topology on the set is a structure that formalizes and specifies a concept called “closeness”. Closeness is defined in a very general way and need not be characterized in terms of a numeric value, metric, or any specific notion of distance. Instead, the topology on *X* is a collection of open sets 𝒯 = {𝒪*_i_*} that satisfy a set of predefined axioms:

1. *X* ∈ 𝒯
2. ∅ ∈ 𝒯
3. All *k* intersections, 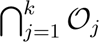, *k* ∈ 1, …, *K* of 𝒯 are in 𝒯
4. All *p* unions 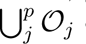 of open sets in 𝒯 are always in 𝒯.

Commonly, topological spaces have the Hausdorff property [27], ensuring separable points in *X*, i.e., for any two points {*x*_1_ ≠ *x*_2_|*x*_1_, *x*_2_ ∈ *X*}, there exist open sets 𝒪_1_ and 𝒪_2_ in 𝒯 such that *x*_1_ ∈ 𝒯_1_, *x*_2_ ∈ 𝒯_2_, and 𝒯_1_ ∩ 𝒯_2_ = ∅.

Although the definition of a topological space permits applicability to highly abstract objects that satisfy the formalism, it is obvious that familiar geometric shapes (without boundaries) in ℝ*^n^* are easily cast as topological metric spaces, with a topology induced by Euclidean or geodesic metrics.

#### 2.4.1. Persistent Homology

Persistent homology, a common method to estimate the topological information from images, point clouds, or graphs, has greatly benefited neuroimaging [28]. Often, persistent homology can be quantified with “persistence diagrams” (PD), examples of which are in figure 2. Persistent homology computes the homological features at different resolutions, a process known as filtration. These features are elements within a given homological class. Homological classes are groupings of mathematical objects based on their properties.

**Figure 2:**
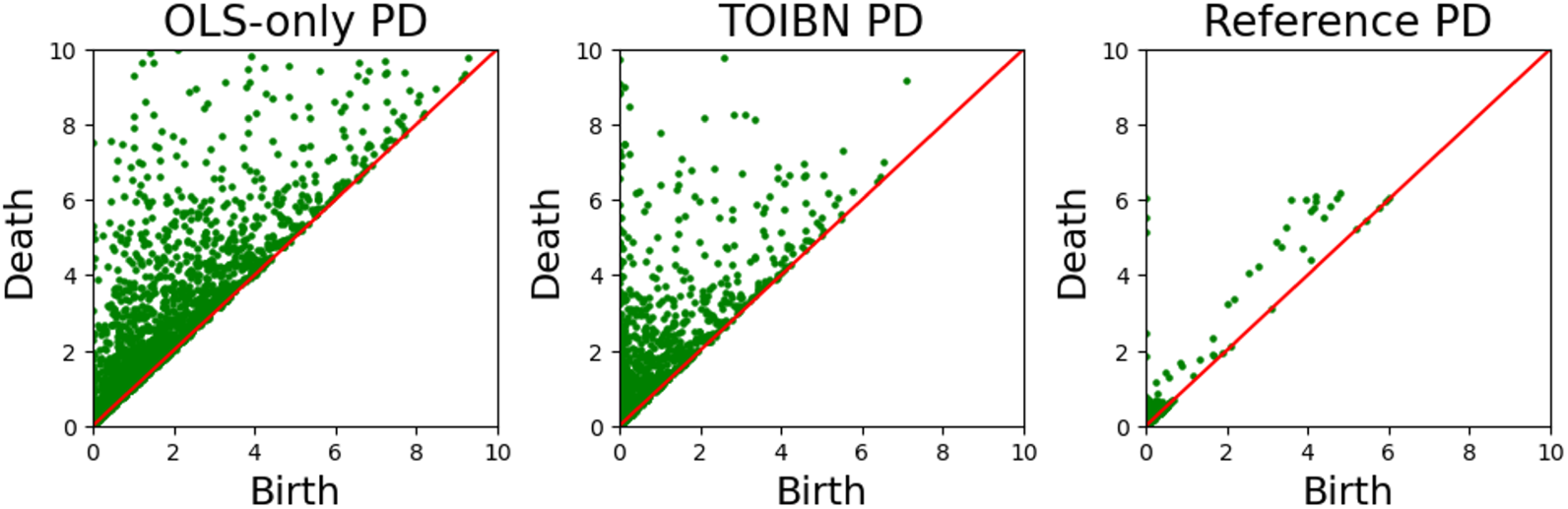
The OLS-only (left), TOIBN (middle), and reference (right) persistence diagrams from the VIS network of one subject. A persistence diagram shows the birth (x-axis) and the death (y-axis) of each homological object (e.g. a component or a hole). In this example, we see that the TOIBN persistence diagram is observably more similar to the reference image than the OLS-only diagram. This is expected, as the loss function is applied directly to these birth-death pairs.

In this work, we use the first two homological classes for topological spaces. H0 and H1, in the 3-dimensional network images. H0 consists of the 0-dimensional connected components, while H1 is the set of holes in the image. As H1 defines 1-dimensional homology in a 3-dimensional space, we refer to this as ”1.5-dimensional” persistence. We omit H2, or the ”hollows”, as estimating this higher homology group is very resource-intensive, to the point of being computationally prohibitive. These homological features are estimated by breaking the image down into simplicial complexes. Each simplicial complex is a set of 0, 1, and 2-dimensional simplices: i.e., points, lines, and triangles. The filtration, specifically level-set filtration, is the set of linearly-spaced resolutions in which the simplices are computed based on a threshold of the voxel intensities. During the persistence computation, we keep track of each H0 and H1 element as they are created or destroyed during the filtration (i.e., a feature exists if it is within the threshold and is destroyed if it is outside the threshold). The creation and destruction of these features are called ”birth” and ”death”. These birth and death values define the overall topological information.

#### 2.4.2. Topology Layer for Machine Learning

The topology layer for machine learning [19], which is the basic part of our algorithm, defines a topological loss function on a given machine learning algorithm learned via gradient descent. One of their primary examples uses this loss function to regularize linear regression. Their method begins by estimating the regression coefficients with ordinary least squares, then iteratively learning the topological loss to produce a final set of coefficients constrained to contain pre-defined topological information. However, the loss function defined in the original paper is ill-fit for our purposes, as it requires “ground-truth” topological information. As this is not available to us, we need to define a loss term that leverages an estimation of the information from the reference maps.

#### 2.4.3. Topological Loss

To properly constrain the spatial maps to include the topological information from the reference maps, a loss function must be defined to compare the subject and reference maps. Here, we leverage the Wasserstein distance [21] to compute the distance between two distributions of homological persistence (i.e., the PDs themselves) [17]. This gives us a convenient minimization problem for the Wasserstein-based loss function over the PD birth/death pairs in Θ*_s_*.

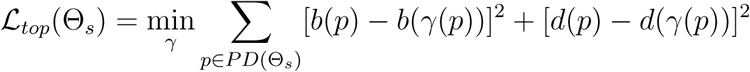

The PDs are computed for both the subject maps, Θ*_s_*, and the reference map Θ*_g_*. *b* and *d* are the births and deaths of the *p^th^*element of the PD, respectively. Each *γ* is a possible bijective mapping between the subject-level and reference-level maps based on the descending sorting of the homological persistences of each map. The optimal *γ* is found by changing the persistence of the subject-level spatial maps (i.e., births/deaths based on voxel intensities), which essentially changes the sort order. And for every object that exists in one spatial map but not the other (i.e., |*PD*(Θ*_s_*)| *>* |*PD*(Θ*_g_*)|), the object is mapped to zero, thus penalizing objects in the estimated subject maps that do not exist in the reference maps. This mapping of erroneous objects to zero is part of this work’s novelty and a vital part of the estimation. Previous work has adapted this loss function to neuroimaging data, but most of it has focused on graphs [17] or brain segmentation [15]. To our knowledge, this is the first use of topological spatial constraints for subject spatial maps.

### 2.5. Our Methodology

Now, we pull these priors together to estimate TOIBN subject-level networks, examples which can be found in figure 3, along with corresponding OLS-only maps. Although the topological layer conceptually works with regression, it requires the “ground truth” topological properties, which we do not have for these reference spatial maps. Thus, we adapt the Wasserstein distance to replace the loss function defined by the machine learning topology layer. This adaptation can be seen in the previous section. As this loss function is differentiable, it can easily be adapted to the topological layer. Although the topological loss does helpfully inform the subject spatial maps, it is only part of the story. We must also include the linear estimation of the subject maps. We implement this linear estimation using the mean squared error (MSE). This gives us the following term:

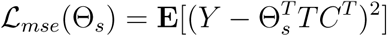

**Figure 3:**
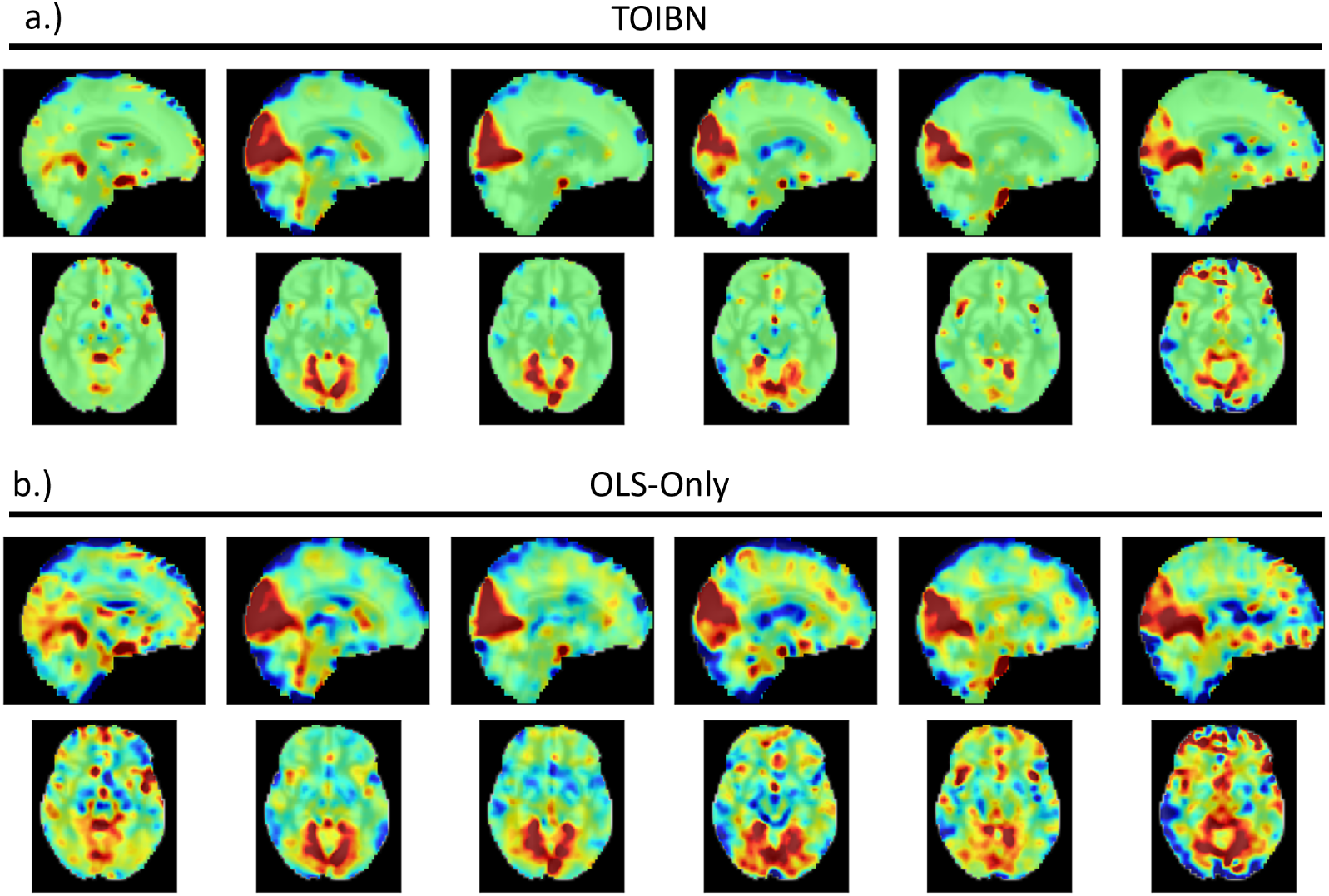
a.) The TOIBN primary VIS maps from six randomly selected subjects, along with the OLS-only maps (b.) from the same subjects.

Where Y is the whole brain fMRI data for the given subject, and *TC* is the timecourse estimated from the reference maps. These *TCs* are the same for both the topological correction and OLS-only methods. It’s also important to note that each term, 𝓛*_top_* and 𝓛*_mse_* has their respective regularization parameter, *λ*, where 0 ≤ *λ* ≤ 1. *λ* is a tunable parameter. For our purposes, we used *λ* = 0.5, so that both loss terms have equal weight. Using both the topological and MSE loss terms, our full training loss function is defined as:

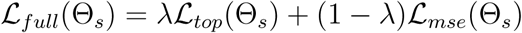

This loss function is then optimized using stochastic gradient descent. As the loss function itself tends to overfit to the topological term, resulting in subject maps that are quite similar to the reference maps, we needed a method for early stopping. Based on empirical evaluations, we found that 150 iterations were sufficient for our purposes. For our method to work properly, the subject spatial map, Θ*_s_* must begin with a sound initial state, to avoid non-optimal local minima during training. This is why we first initialize every subject spatial map as the OLS best fit using the Moore-Penrose pseudoinverse. Importantly, this OLS is computed using all network timecourses, but the topological correction algorithm is computed per network. A diagram of our methodology is in figure 4.

**Figure 4:**
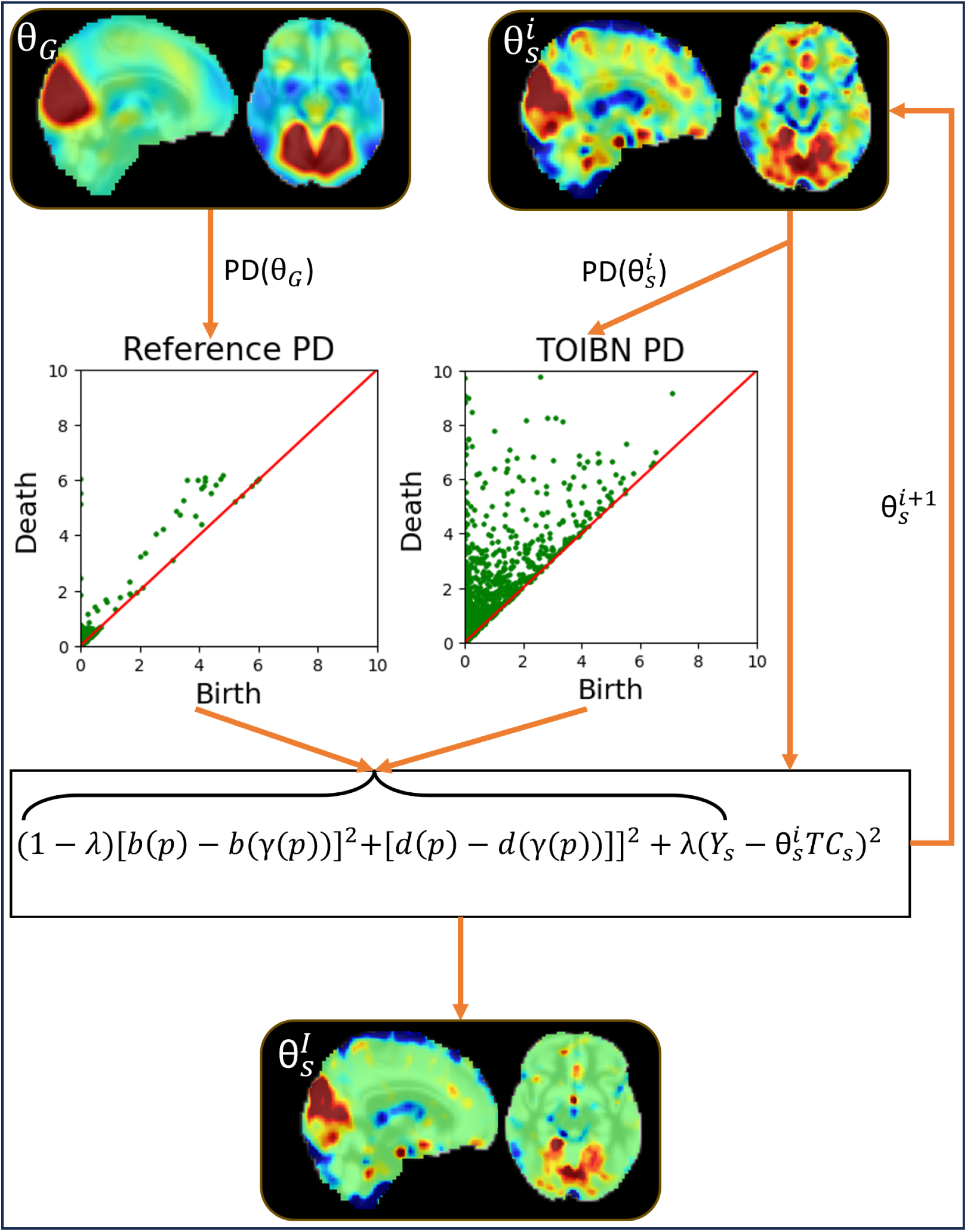
A diagram of our methodology. We start by estimating the reference network, *θ_G_*, and its associated persistence diagram, *PD*(*θ_G_*). Then, we initialize the subject network, such that 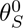 is the OLS estimation. For each iteration *i*, we compute the subject PD, 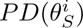 and estimate 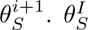 is the final subject estimation.

### 2.6. Training and Gradients

The gradients for the training phase are computed the same way as laid out in the original paper [19]. Essentially, each birth-death pair can be mapped to a single voxel. For the 2D case, as laid out in their paper, every homological object is mapped to a single simplex, or triangle, line, and point, which is then mapped to a single voxel in the subject maps (Θ*_s_*), such that each voxel maps to one point, two lines, and two triangles; excluding edge voxels. However, in our case, every voxel now maps to points, lines, and triangles in 3 dimensions. This means that each voxel maps to one point, three lines, and two triangles. For the gradient computation, a gradient is computed for each homological object, and this gradient is then applied to a given voxel based on this mapping.

### 2.7. Experiments

To show the validity of our method, we suggest that there are three over-arching empirical goals: show that the TOIBN maps are more similar to the reference maps than the OLS-only maps, the subject variability of the spatial maps is better retained by the topological correction maps, and finally that the topological correction aids the image quality. We evaluate the first by comparing the subject maps with the reference maps in two ways, voxel-wise similarity measures and topological similarities. Since the persistences between the reference maps and the TOIBN maps are more similar than the OLS-only maps and the reference maps, we compare their topological similarity using measurements that quantify the homological objects in a way that aggregates the homological properties, which is juxtaposed against persistence, a measure that is specific to the homological dimensions. After binarizing the maps based on the reference-level maps, we compare the Euler characteristic for all subject maps and the reference map for every network and statistically compare the subject maps and the reference maps for both the OLS-only and TOIBN maps. The Euler characteristic is defined for topological properties simply as the number of connected components - the number of holes + the number of hollows. After verifying that the TOIBN maps are more topologically similar to the reference than the OLS-only maps, we measure and quantify the geometric differences between the reference maps and the subject maps. We will accomplish this with the mean absolute error (MAE) as a distance function between the reference maps and subject maps (both OLS-only and TOIBN).

To test the efficacy of the topological correction toward the second goal, we lay out two sets of experiments. The first is to test how similar the TOIBN subjects are to the reference maps compared to the OLS-only subjects. Here, we use Pearson correlation between pairs of subject spatial maps as a measure of similarity, or how much the spatial maps vary from subject to subject. In this same vein, we analyze not only the full subject variability but also the voxel-wise variability based on the voxels that are most relevant to the reference-level map. We hypothesize that the TOIBN maps have more subject variance only in voxels that are relevant to the reference map. We do this by testing the subject standard deviation of the voxels with the highest values in the reference maps. Because of sign ambiguity, the highest values and the most negative values can both be considered relevant, however, it is generally accepted that the highest values are the voxels that make up the network. We found these voxels by clustering every voxel in a given reference network map using k-means. Our k-means model clusters each spatial map into three clusters: low-valued, middle-valued, and high-valued. We chose three because we essentially view these voxel values as three categories: the highest values most relevant to the network, the values closest to zero, and the most negative values.

Next, we want to show that the TOIBN maps have important and interpretable impacts on the maps as a whole. Namely, we show that the maps are essentially cleaner, as estimated by the contrast-to-noise ratio (CNR). Historically, subject-level brain network estimations have relatively low CNR [29]. This is due primarily to the low CNR of the BOLD fMRI signal itself. If topological correction is to be a viable addition for subject-level network estimation, then it must improve the image quality in some capacity.

CNR, generally defined as (*µ*(*in*)−*µ*(*out*))/*σ*(*noise*), is an oft-discussed measurement with many possibilities for estimation. We chose to estimate the *in* and *out* portions using the same K-Means clusters from the previous analyses. Although there is no perfect “ground-truth” contrast, we can use an estimator that can separate the highest value points (assumed to be most relevant to the network) from the rest of the map. Using k-means we found three total clusters, again to represent high values, middle values, and low values. Using the samples from the highest-valued cluster as the *in* values, and the samples from the center cluster as the *out* values, we computed (*µ*(*in*) − *µ*(*out*)). Finally, to estimate the standard deviation of the noise, we consider the cerebral-spinal fluid (CSF) of the ventricles, as it does not contain neuronally related signals. This gives us a final equation of (*µ*(*highest cluster*) − *µ*(*center voxels*))/*σ*(*CSF* ). As a final note, k-means is only performed on voxels that are not considered CSF.

Finally, we analyze how the topological properties differ between individuals with schizophrenia and controls. We compare these group differences between the OLS-only and topologically-corrected maps. We do this by statistically comparing the aforementioned Euler characteristics between patients with schizophrenia and controls.

## 3. Results

### 3.1. Topological Similarity to the Reference Maps

We found, as expected, that the TOIBN maps are indeed more similar to the reference maps than the OLS-only maps. The Euler characteristic is a different estimation of topological properties than homological persistence. Figure 5 (top) shows the results of this analysis, and how the topological correction impacts the OLS-only maps. This plot is the distribution of the differences between the reference maps and the TOIBN maps (red) compared to the differences between the OLS-only and reference maps (green) for all 14 networks.

**Figure 5:**
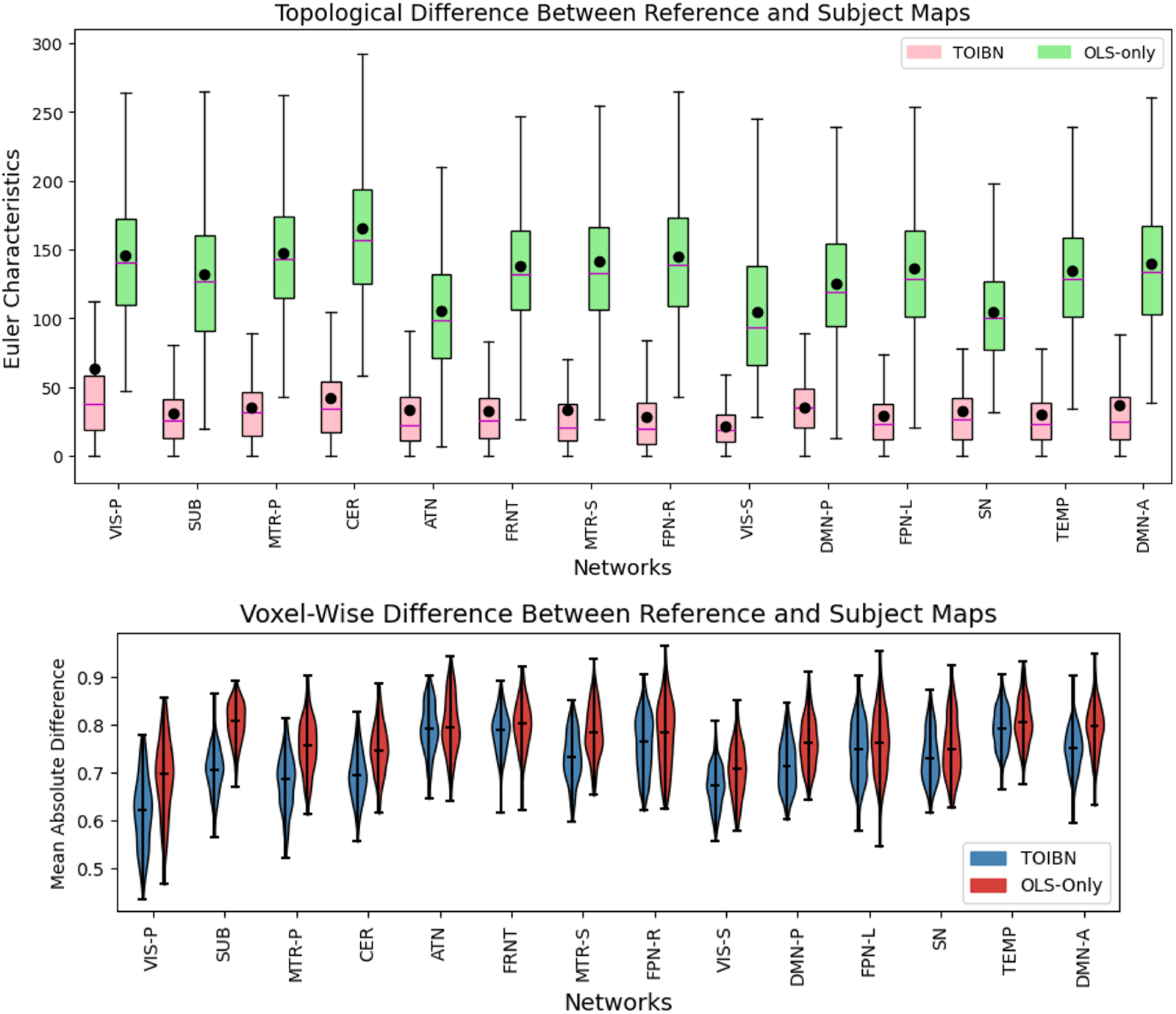
Top) The differences between the per-subject Euler characteristic of the TOIBN maps and the Euler characteristic of the reference image (pink) compared to the difference between the reference map and the OLS-only maps (green). The solid purple lines are the medians, and the black circles are the means. Compared to the Euler characteristics between the OLS-only and reference maps. This Euler characteristic is defined on a binarized image, where every subject is, after z-scoring, binarized at a threshold of 0. As expected, the TOIBN maps are more similar to the reference maps than the OLS-only maps. Bottom) Plots of the per-subject distance (defined as the MAE) to the reference image for all TOIBN subject maps (blue) and all OLS-only maps (red). So, not only are the maps more topologically similar to the reference maps, but they are also statistically similar. All networks marked with an asterisk are statistically significant after FDR correction.

### 3.2. Voxel-wise Similarity to the Reference Maps

To define the distance between the subject and reference maps, we use the MAE between the two maps. Figure 5 (bottom) shows the distribution of per-voxel MAEs between every TOIBN subject map and OLS-only map for each network. Table 1 shows a table of the statistical tests between the TOIBN MAEs and the OLS-only MAEs. Overall, we see that the topological correction does indeed accomplish our first goal. The corrected maps are statistically much more similar to the reference maps using inferred information without direct and forceful constraints.

**Table 1:** Statistical differences between the reference-subject similarities of TOIBN maps and OLS-only. After FDR correction, asterisks denote networks are statistically significant, with all P-Values < 0.001. As such, the topologically-corrected maps are significantly more similar to the reference maps than the OLS-only maps.

|  |  |  |  |  |  |  |  |
| --- | --- | --- | --- | --- | --- | --- | --- |
|  | VIS-P | SUB | MTR-P | CER | ATN | FRNT | MTR-S |
| T-Value | *71.35 | *96.20 | *81.00 | *55.99 | *5.29 | *16.02 | *60.88 |
|  | FPN-R | VIS-S | DMN-P | FPN-L | SN | TEMP | DMN-A |
| T-Value | *20.95 | *42.34 | *60.56 | *20.30 | *23.45 | *17.47 | *50.72 |

### 3.3. Subject Variability

Figure 6 shows the results of our subject variability analyses. The top image compares the two distributions (TOIBN and OLS-only) of the between-subject Pearson correlations. Here we very clearly see that the TOIBN subjects are much less correlated, or more dissimilar than the OLS-only maps. Table 2 shows that the TOIBN maps are significantly less correlated than the OLS-only maps. Thus, we suggest that the topological correction promotes subject variability.

**Figure 6:**
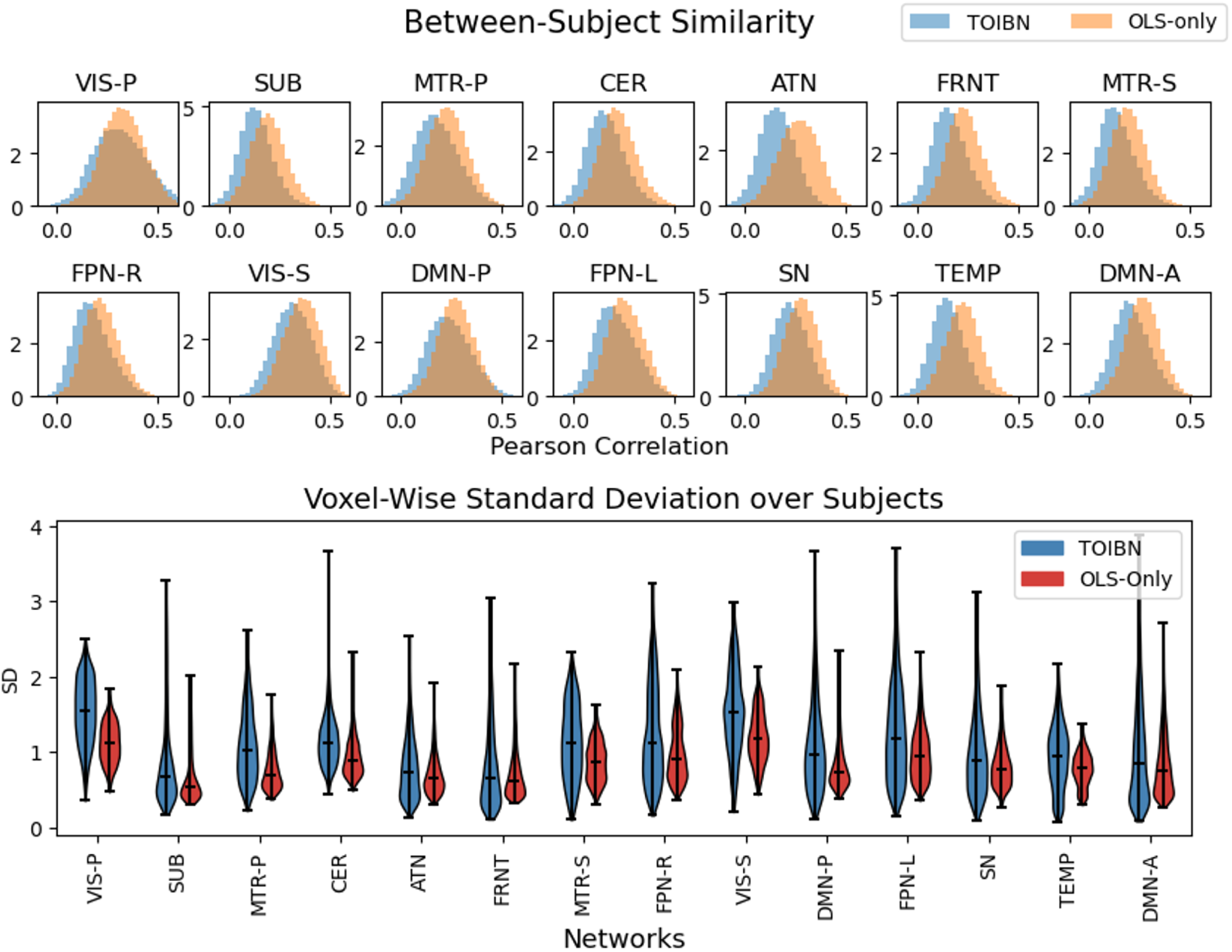
Top) Histograms of the pair-wise subject spatial correlation (Pearson correlation). This figure shows that, for every network, the TOIBN maps are less correlated, implying that there is higher subject variation. Each bin is normalized as counts / (sum(counts) * diff(bins)). These correlations are how we begin to estimate the subject-specificity of the methods, by capturing the subject variability. Bottom) Plots of the per-voxel subject standard deviation (SD) for all TOIBN subject maps (blue) and all OLS-only maps (red) for voxels that are most positively relevant for the reference maps. We find these highly-correlated voxels by clustering every reference image into three clusters, then selecting the cluster with the highest value to be the correlated value, and removing the remaining voxels. From this, we see that the standard deviation over the subjects is higher for the TOIBN maps for voxels that are inside the network. This complements the correlation results, showing that there is more subject variability from the TOIBN maps, but that this variability exists inside the networks. In both figures, all networks are statistically significantly different after FDR correction.

**Table 2:** Statistical differences among the between-subject similarities of TOIBN maps and OLS-only. From these, we see the statistical difference between the correlation distributions, showing that the TOIBN maps are significantly less correlated than the OLS-only maps. After FDR correction, all networks with asterisks are statistically significant, with all P-Values < .001.

|  |  |  |  |  |  |  |  |
| --- | --- | --- | --- | --- | --- | --- | --- |
|  | VIS-P | SUB | MTR-P | CER | ATN | FRNT | MTR-S |
| T-Value | *101.80 | *376.38 | *354.90 | *470.13 | *667.40 | *518.26 | *459.05 |
|  | FPN-R | VIS-S | DMN-P | FPN-L | SN | TEMP | DMN-A |
| T-Value | *450.05 | *494.62 | *218.99 | *439.51 | *473.45 | *577.98 | *350.02 |

Although this is a promising start, we also show that the subject variability is highest in regions that are most related to the network. The bottom image of figure 6 shows that even when restricted to the network-relevant voxels, subject variability does indeed increase after topological correction. Table 3 shows the statistical tests between the TOIBN and OLS-only maps.

**Table 3:** Statistical differences using the Kruskal-Wallis test, between the relevant-voxel-wise standard deviations of TOIBN maps and OLS-only. From these, we see the statistical difference between the standard deviation distributions within the most relevant areas, showing that the TOIBN maps are significantly less correlated than the OLS-only maps. These relevant areas are defined using K-Means clustering for each subject. Kruskal-Wallis was chosen due to the non-Gaussian nature of standard deviations. After FDR correction, all networks, except the FRNT, are statistically significant (denoted with an asterisk), with P-Values < .001.

|  | VIS-P | SUB | MTR-P | CER | ATN | FRNT | MTR-S |
| --- | --- | --- | --- | --- | --- | --- | --- |
| T-Value | *1204.50 | *145.67 | *876.70 | *1619.25 | *85.28 | 1.49 | *691.30 |
|  | FPN-R | VIS-S | DMN-P | FPN-L | SN | TEMP | DMN-A |
| T-Value | *350.08 | *944.80 | *401.92 | *388.24 | *161.96 | *512.88 | *35.09 |

### 3.4. Contrast-to-Noise Ratio

The final piece of our puzzle is the image quality after topological correction. Figure 7 shows the distributions of the CNR for each TOIBN (blue) map compared to the OLS-only (orange) maps. Table 4 shows the statistical tests between these distributions. We see that the TOIBN maps have a statistically higher CNR, suggesting that the topological correction reduces the overall noise.

**Figure 7:**
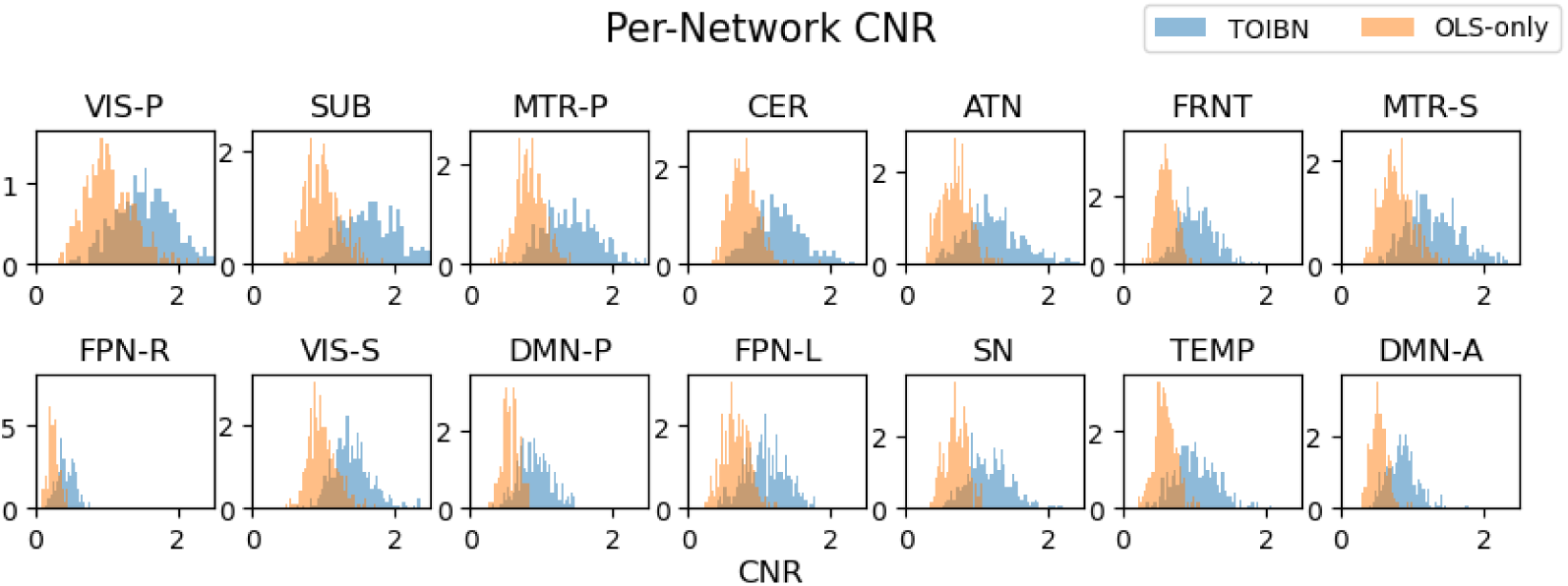
Histograms of the per-subject CNRs for each network. The CNR is computed as the average of the voxels in the network minus the average of the voxels outside of the network, divided by the SD of the noise, which we define as the CSF. The voxels inside and outside of the network are defined using K-Means clustering. We cluster every image into three clusters and select the cluster with the highest value to be the network values and every voxel from the other two clusters to be the values outside of the network. We see that the CNR of the TOIBN maps is significantly (see table 4) lower than the OLS-only CNR values.

**Table 4:** Statistical differences between the CNR from the topologically-corrected and OLS-only maps. We see that, for every network, the TOIBN maps have significantly higher CNRs. After FDR correction, all networks are statistically significant, with all P-Values < .001.

| VIS-P | SUB | MTR-P | CER | ATN | FRNT | MTR-S |
| --- | --- | --- | --- | --- | --- | --- |
| *47.63 | *50.45 | *45.66 | *39.72 | *44.20 | *49.68 | *49.91 |
| FPN-R | VIS-S | DMN-P | FPN-L | SN | TEMP | DMN-A |
| *49.28 | *51.65 | *49.80 | *50.31 | *48.54 | *44.13 | *47.23 |

### 3.5. Group differences

We show that the topological properties of the TOIBN maps reveal more group differences between patients and controls. We computed the group differences using unpaired, two-sample t-tests and then corrected the p-values with the false discovery rate (FDR) correction for multiple comparisons. Figure 8 illustrates these differences. Overall, see more networks that are significantly different between groups among the TOIBN maps than the OLS-only maps.

**Figure 8:**
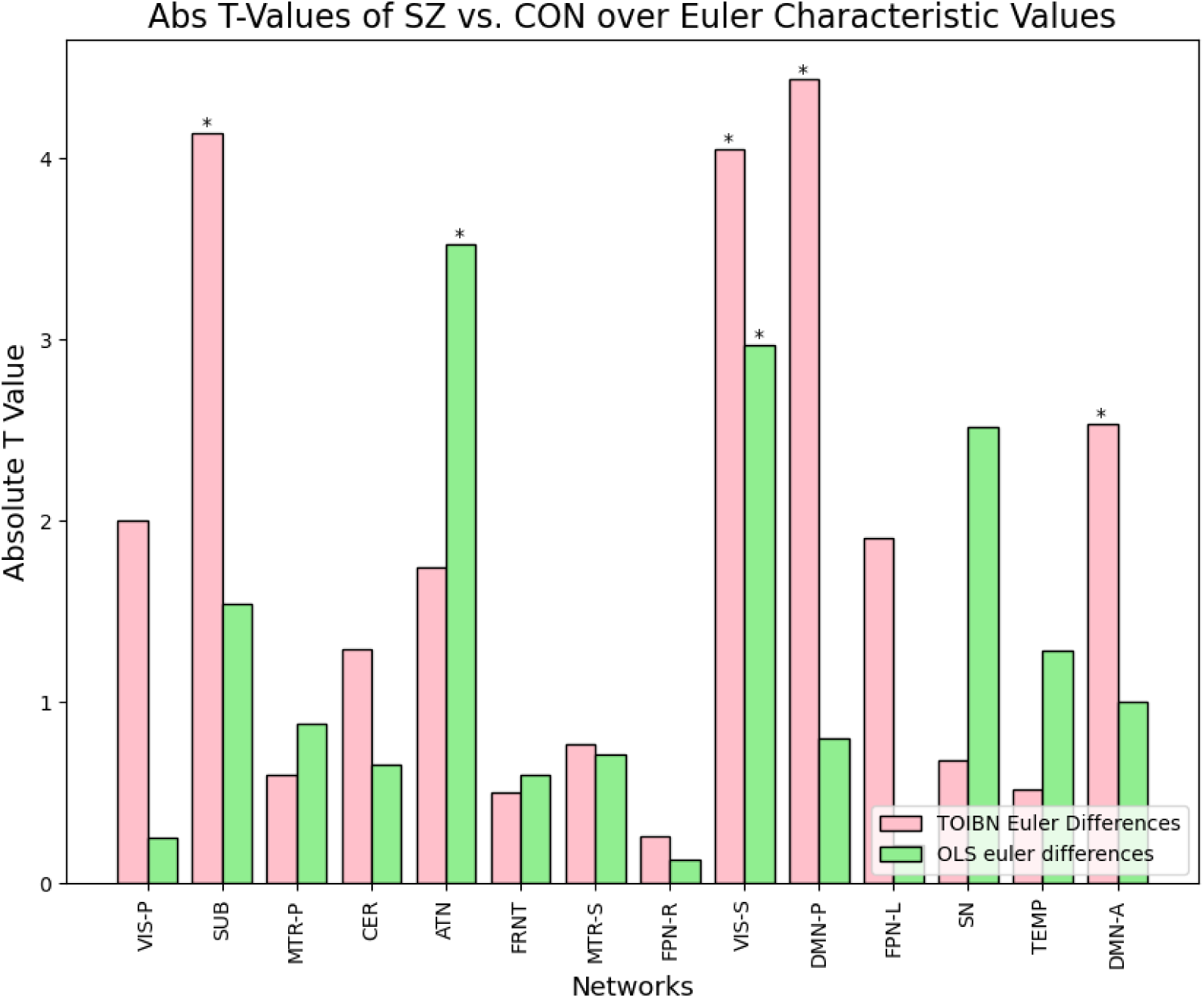
Group differences of the Euler characteristics between patients and controls for the TOIBN maps (pink) and OLS-only maps (green). The stars indicate which networks are significantly different between groups using FDR-corrected, unpaired, two-sample t-tests. The SUB, secondary VIS, posterior DMN, And anterior DMN networks of the TOIBN maps are significantly different between groups. Whereas only the ATN and secondary VIS are significantly different among the OLS-only maps.

## 4. Discussion

The past several decades have seen prolific use of functional brain networks estimated from fMRI data. These networks, estimated in many different ways, have been invaluable for understanding how brain function works, both at rest and during task-based experiments. Often, the field estimates networks from groups of subjects. These group networks, or reference networks, are often very effective at capturing spatial properties of human brain function. However, this estimation becomes more difficult at the subject level. This work is a first step towards subject-level spatial map estimation that leverages tertiary reference-level information as a soft constraint. Our preliminary results show that this methodology results in subject maps that are more similar to the reference maps while enhancing subject variation and increasing the CNR, which can often be a problem for subject maps. Beyond this, the TOIBN maps have topologies that are generally more sensitive to group differences between patients and controls.

This work is only the beginning of what is possible for topologically constrained subject spatial maps. There is a wealth of future work that is possible. Primarily, the next steps, with better computational resources or more time, should be to estimate the 2*^nd^* dimensional homological objects, or the hollows, and compute the PDs for these objects as well as the 0D and 1D objects. This should provide a much better assessment of the topological properties of the 3D spatial maps. This lack of 2D homological object estimation is a primary limitation of the current work. Another major limitation is the reliance on an empirically chosen number of iterations, instead of a clear and well-thought-out stopping criteria. Future work will include research and experimentation into possible stopping criteria. Primarily, we will experiment with backtracking to find a proper stopping point. Finally, due to the scope of this project, we could not reasonably include comparisons between the TOIBN method and recent spatially constrained methods. These spatially-constrained methods are very powerful, and we would be remiss not to compare their efficacy to our soft-constrained method in the future.

Overall, we think that these preliminary results validate topological information as a useful soft constraint on subject-level network estimation. This now opens the door to a host of possibilities. After developing new avenues with STR-specific subject estimation, we can adapt this methodology to other estimation methods. Promisingly, we can adapt this methodology to subject-level ICA, directly computing each subject’s networks while imposing this soft constraint.

## 5. Remarks and Declarations

- All code will be made available upon request or publication.
- We have no competing or conflicting interests in the publication of this manuscript.
- This is an original work that has not been published elsewhere and it is not currently being considered for publication elsewhere.
- This work was supported by the National Institutes of Health grants R01MH123610 and R01MH118695, as well as the National Science Foundation grant 2112455 (to Calhoun VD).

## 6. Author Contributions

- Dr. Noah Lewis developed the method, implemented the code, compiled the results, and wrote the paper
- Dr. Armin Iraji assisted in analyzing and quantifying the resulting subject networks from the method, as well as editing the paper
- Dr. Robyn Miller assisted with method development
- Dr. Oktay Agcaoglu helped quantify the results and edit the paper
- Dr. Vince Calhoun advised the entire project, assisted with ideation and editing the paper, as well as funding acquisition

## Notes

### Competing Interest Statement

The authors have declared no competing interest.

